# Mosaic nanoparticles elicit cross-reactive immune responses to zoonotic coronaviruses in mice

**DOI:** 10.1101/2020.11.17.387092

**Authors:** Alexander A. Cohen, Priyanthi N.P. Gnanapragasam, Yu E. Lee, Pauline R. Hoffman, Susan Ou, Leesa M. Kakutani, Jennifer R. Keeffe, Hung-Jen Wu, Mark Howarth, Anthony P. West, Christopher O. Barnes, Michel C. Nussenzweig, Pamela J. Bjorkman

## Abstract

Protection against SARS-CoV-2 and SARS-related emergent zoonotic coronaviruses is urgently needed. We made homotypic nanoparticles displaying the receptor-binding domain (RBD) of SARS-CoV-2 or co-displaying SARS-CoV-2 RBD along with RBDs from animal betacoronaviruses that represent threats to humans (mosaic nanoparticles; 4-8 distinct RBDs). Mice immunized with RBD-nanoparticles, but not soluble antigen, elicited cross-reactive binding and neutralization responses. Mosaic-RBD-nanoparticles elicited antibodies with superior cross-reactive recognition of heterologous RBDs compared to sera from immunizations with homotypic SARS-CoV-2–RBD-nanoparticles or COVID-19 convalescent human plasmas. Moreover, sera from mosaic-RBD–immunized mice neutralized heterologous pseudotyped coronaviruses equivalently or better after priming than sera from homotypic SARS-CoV-2–RBD-nanoparticle immunizations, demonstrating no immunogenicity loss against particular RBDs resulting from co-display. A single immunization with mosaic-RBD-nanoparticles provides a potential strategy to simultaneously protect against SARS-CoV-2 and emerging zoonotic coronaviruses.

**One sentence summary:** Nanoparticle strategy for pan-sarbecovirus vaccine

**125-character summary for online ToC:** Immunizing with nanoparticles displaying diverse coronavirus RBDs elicits cross-reactive and neutralizing antibody responses.

## Main Text

SARS-CoV-2, a newly-emergent betacoronavirus, resulted in a global pandemic in 2020, infecting millions and causing the respiratory disease COVID-19 (*1*, *2*). Two other zoonotic betacoronaviruses, SARS-CoV and MERS-CoV, also resulted in outbreaks within the last 20 years (*3*). All three viruses presumably originated in bats (*4*), with SARS-CoV and MERS-CoV adapting to intermediary animal hosts before jumping to humans. SARS-like viruses circulate in bats and serological surveillance of people living near caves where bats carry diverse coronaviruses demonstrates direct transmission of SARS-like viruses with pandemic potential (*5*), suggesting a pan-coronavirus vaccine is needed to protect against future outbreaks and pandemics. In particular, the bat WIV1 and SHC014 strains are thought to represent an ongoing threat to humans (*6*, *7*).

Most current SARS-CoV-2 vaccine candidates include the spike trimer (S), the viral protein that mediates target cell entry after one or more of its receptor-binding domains (RBDs) adopt an “up” position to bind a host receptor (Fig. 1A). The RBDs of human coronaviruses SARS-CoV-2, SARS-CoV, HCoV-NL63, and related animal coronaviruses (WIV1 and SCH014) use angiotensin-converting enzyme 2 (ACE2) as their host receptor (*1, 8, 9*), while other coronaviruses use receptors such as dipeptidyl peptidase 4 (*10*) or sialic acids (*11*, *12*). Consistent with its function in viral entry, S is the primary target of neutralizing antibodies (*13–22*), with many targeting the RBD (*14–18, 21–26*).

**Figure 1.**
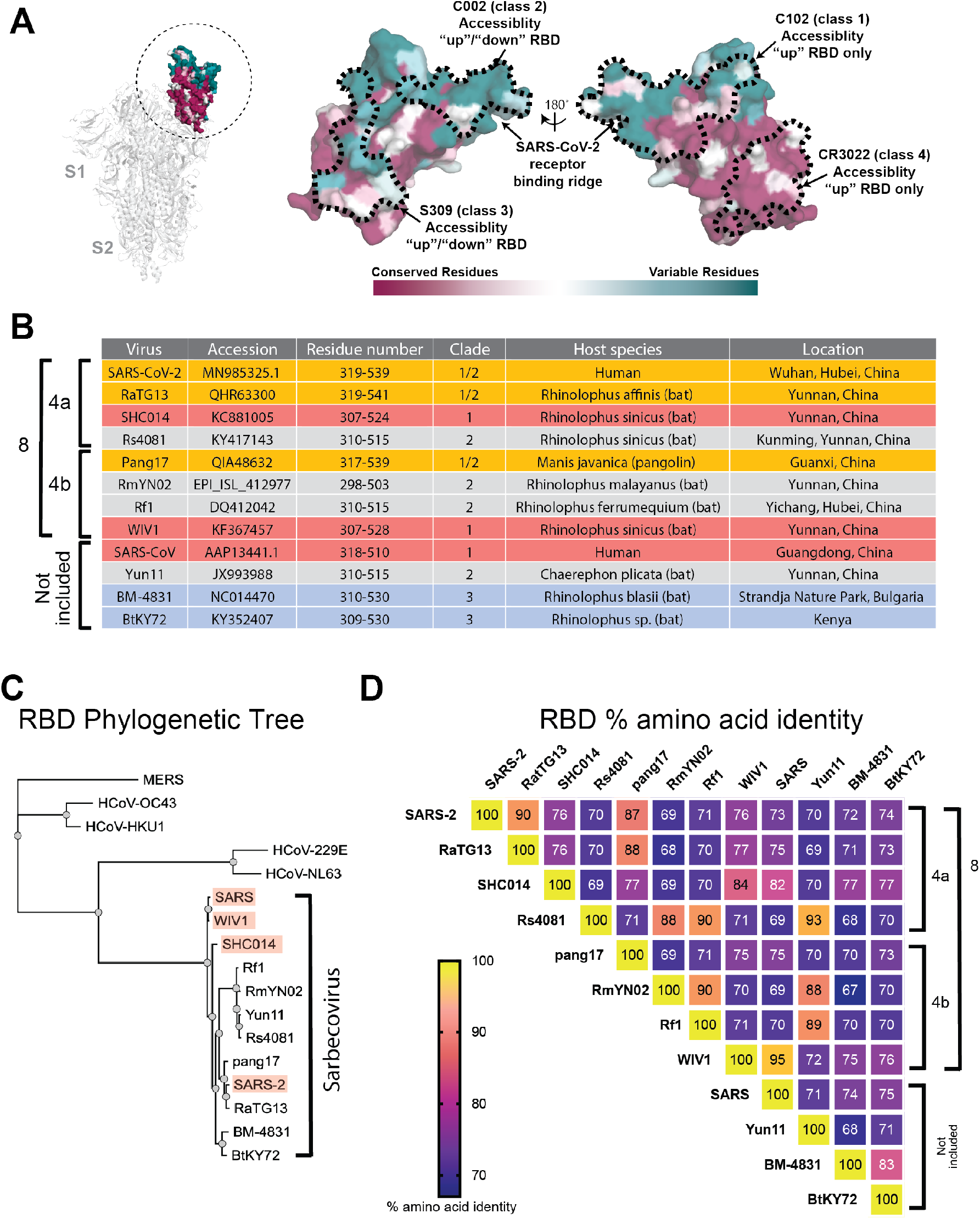
Properties of RBDs chosen for this study. (A) Left: Structure of SARS-CoV-2 S trimer (PDB 6VXX) with one RBD (dashed circle) in an “up” position. Middle and right: Sequence conservation of 12 RBDs calculated by the ConSurf Database (*49*) plotted on a surface representation of the RBD structure (PDB 7BZ5). Epitopes for representatives from defined classes of RBD-binding antibodies (class 1-class 4) (*24*) indicated by dashed lines. (B) Summary of properties of the viral strains from which the 12 sarbecovirus RBDs were derived. (C) Phylogenetic tree of human and selected other coronaviruses based on RBD protein sequences. Red shading indicates strains known to use ACE2 as a receptor. (D) Heat map showing percent amino acid sequence identities between 12 sarbecovirus RBDs.

Multivalent display of antigen enhances B-cell responses and can provide longer-lasting immunity than monovalent antigens (*27*, *28*), thus protein-based vaccine candidates often involve a nanoparticle that enables antigen multimerization. Many nanoparticles and coupling strategies have been explored for vaccine design (*29*), with “plug and display” strategies being especially useful (*30*, *31*). In one such approach, multiple copies of an engineered protein domain called SpyCatcher fused to subunits of a virus-like particle form spontaneous isopeptide bonds to purified antigens tagged with a 13-residue SpyTag (*29*–*32*). The SpyCatcher-SpyTag system was used to prepare multimerized SARS-CoV-2 RBD or S trimer that elicited high titers of neutralizing antibodies (*33*, *34*). Although promising for protection against SARS-CoV-2, coronavirus reservoirs in bats suggest future cross-species transmission (*6, 7, 35*), necessitating a vaccine that protects against emerging coronaviruses as well as SARS-CoV-2. Here we prepared SpyCatcher003-mi3 nanoparticles (*31*, *36*) simultaneously displaying SpyTagged RBDs from human and animal coronaviruses to evaluate whether mosaic particles can elicit cross-reactive antibody responses, as previously demonstrated for influenza head domain mosaic particles (*37*). We show that mice immunized with homotypic or mosaic nanoparticles produced broad binding and neutralizing responses, in contrast to plasma antibodies elicited in humans by SARS-CoV-2 infection. Moreover, mosaic nanoparticles showed enhanced heterologous binding and neutralization properties against human and bat SARS-like betacoronaviruses (sarbecoviruses) compared with homotypic SARS-CoV-2 nanoparticles.

We used a study of sarbecovirus RBD receptor usage and cell tropism (*38*) to guide our choice of RBDs for co-display on mosaic particles. From 29 RBDs that were classified into distinct clades (clades 1, 2, 1/2, and 3) (*38*), we identified diverse RBDs from SARS-CoV, WIV1, and SHC014 (clade 1), SARS-CoV-2 (clade 1/2), Rs4081, Yunnan 2011 (Yun11), and Rf1 (clade 2), and BM48-31 (clade 3), of which SARS-CoV-2 and SARS-CoV are human coronaviruses and the rest are bat viruses originating in China or Bulgaria (BM48-31). We also included RBDs from the GX pangolin clade 1/2 coronavirus (referred to here as pang17) (*39*), RaTG13, the bat clade 1/2 virus most closely related to SARS-CoV-2 (*40*), RmYN02, a clade 2 bat virus from China (*41*), and BtKY72, a Kenyan bat clade 3 virus (*42*). Mapping of the sequence conservation across selected RBDs showed varying degrees of sequence identity (68-95%), with highest sequence variability in residues corresponding to the SARS-CoV-2 ACE2 receptor-binding motif (Fig. 1A-D; fig. S1). We chose 8 of the 12 RBDs for making three types of mosaic nanoparticles: mosaic-4a (coupled to SARS-2, RaTG13, SHC014, and Rs4081 RBDs), mosaic-4b (coupled to pang17, RmYN02, RF1, and WIV1 RBDs), and mosaic-8 (coupled to all eight RBDs), and compared them with homotypic mi3 particles constructed from SARS-CoV-2 RBD alone (homotypic SARS-2). RBDs from SARS, Yun11, BM-4831, and BtKY72, which were not coupled to mosaic particles, were used to evaluate sera for cross-reactive responses.

SpyTag003-RBDs were coupled to SpyCatcher003-mi3 (60 potential conjugation sites) (*36*, *43*) to make homotypic and mosaic nanoparticles (Fig 2A). Particles were purified by size exclusion chromatography (SEC) and analyzed by SDS-PAGE, revealing monodisperse SEC profiles and nearly 100% conjugation (Fig. 2B,C). Representative RBDs were conjugated to SpyCatcher003-mi3 with similar or identical efficiencies (fig. S2), suggesting that mosaic particles contained approximately equimolar mixtures of different RBDs.

**Figure 2.**
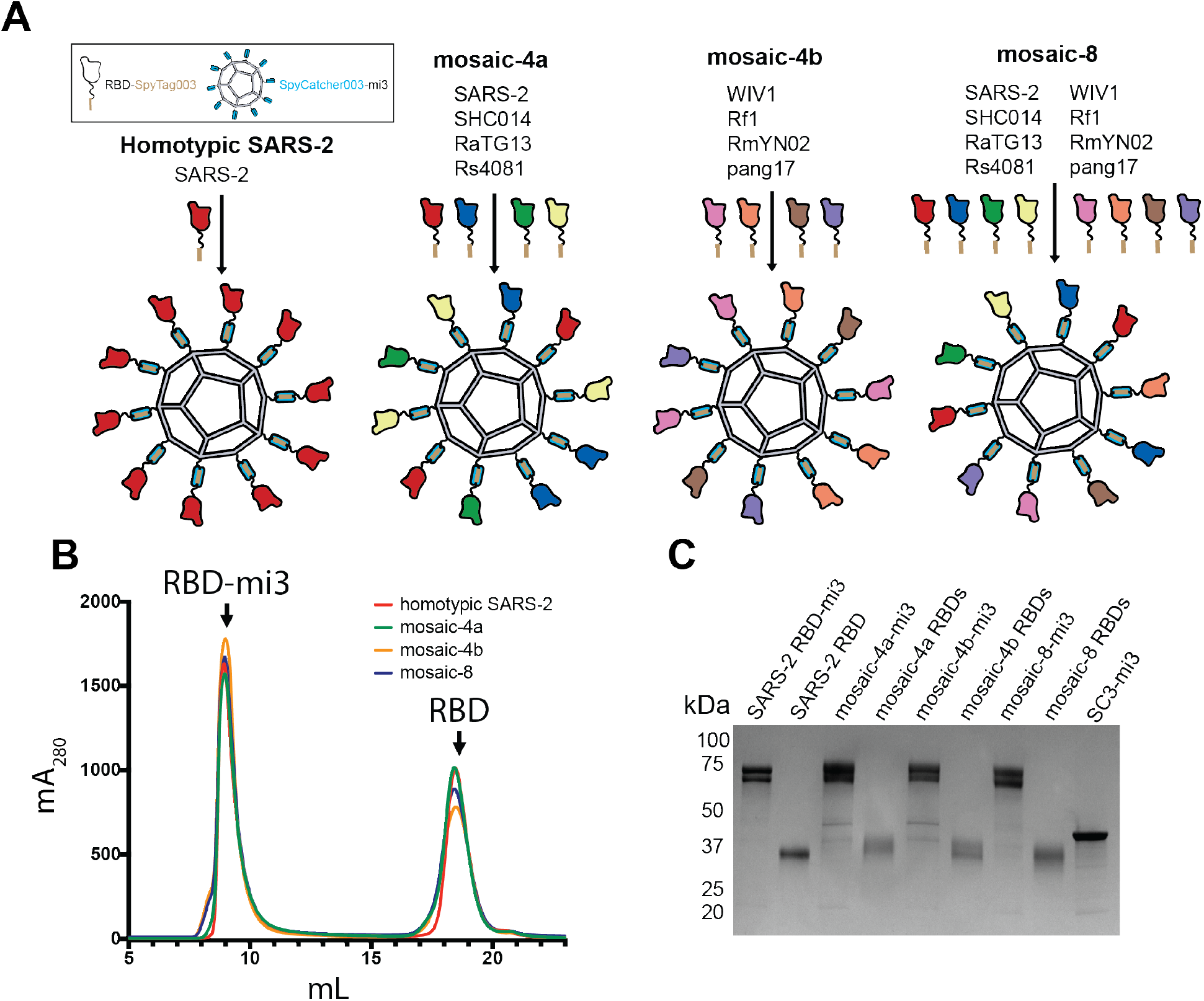
Construction of RBD nanoparticles. (A) Left: SpyTagged RBDs were attached to SpyCatcher003-mi3 to make a homotypic particle and three mosaic particles. 10 of 60 potential coupling sites on mi3 are shown for clarity. (B) SEC profile showing separation of RBD nanoparticles and free RBD proteins. (C) Coomassie-stained SDS-PAGE of RBD-coupled nanoparticles, free RBD proteins, and uncoupled SpyCatcher003-mi3 particles (SC3-mi3).

We immunized mice with either soluble SARS-CoV-2 spike trimer (SARS-2 S), nanoparticles displaying only SARS-2 RBD (homotypic SARS-2), nanoparticles co-displaying RBDs (mosaic-4a, mosaic-4b, mosaic-8), or unconjugated nanoparticles (mi3). IgG responses were evaluated after prime or boost immunizations (Fig. 3A) by ELISA against SARS-2 S (Fig. 3B) or a panel of RBDs (Fig. 3C-F; fig. S3). Sera from unconjugated nanoparticle-immunized animals (black in Fig. 3, fig. S3) showed no responses above background. Anti-SARS-2 S trimer and anti-SARS-2 RBD serum responses were similar (Fig. 3B,C), demonstrating that antibodies elicited against RBDs can access their epitopes on SARS-2 S trimer. We also conducted in vitro neutralization assays using a pseudotyped virus assay that quantitatively correlates with authentic virus neutralization (*44*) for strains known to infect 293T_ACE2_ target cells (SARS-CoV-2, SARS, WIV1 and SHC104). Neutralization and ELISA titers were significantly correlated (fig. S4), thus suggesting ELISAs are predictive of neutralization results when pseudotyped neutralization assays were not possible due to unknown viral entry receptor usage.

**Figure 3.**
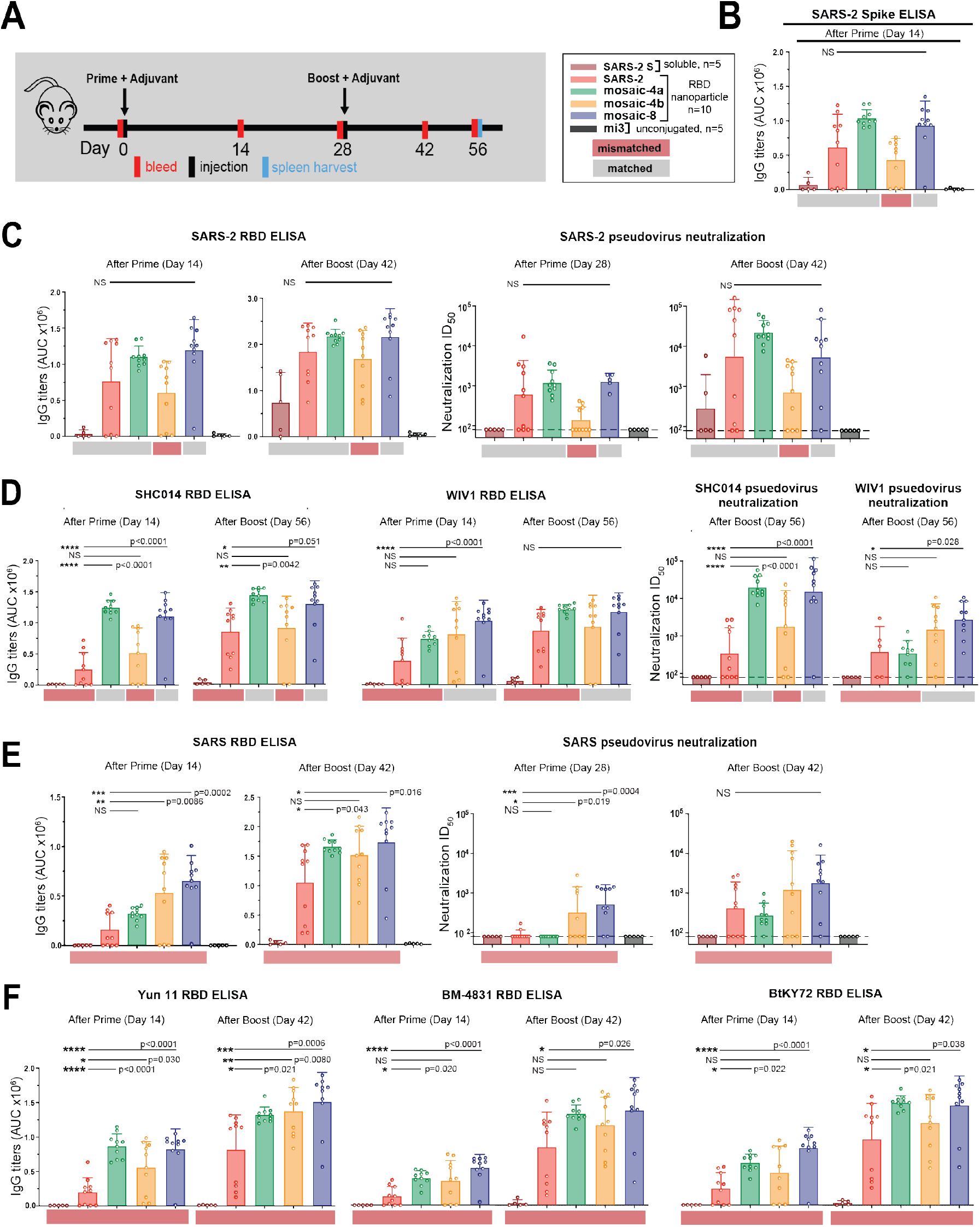
RBD nanoparticles induce cross-reactive IgG responses in immunized mice. Red and gray rectangles below ELISA and neutralization data represent mismatched strains (red; RBD from that strain was not present on the immunized particle) or matched strains (gray; RBD was present on the immunized particle). (A) Left: Immunization schedule. Adjuvant=AddaVax (Invivogen). Right: Key for immunizations; number of mice in each cohort is indicated. (B-F) Mice were immunized with soluble SARS-CoV-2 S trimer (SARS-2 S; brown bars), or the following nanoparticles: homotypic SARS-2 (red), mosaic-4a (green), mosaic-4b (yellow), mosaic-8 (blue), or unconjugated SpyCatcher003-mi3 (mi3; black). ELISA data from serum IgG responses to SARS-2 spike trimer (B) or RBDs (C-F) shown as area under the curve (AUC). For C-E, neutralization potencies are presented as half-maximal inhibitory dilutions (ID_50_ values) of sera against the pseudoviruses from the indicated coronavirus strains. Dashed horizontal lines correspond to the lowest dilution representing the limit of detection. Each dot represents serum from one animal, with means and standard deviations for vaccinated cohorts represented by rectangles (mean) and horizontal lines (SD). Significant differences between groups linked by horizontal lines are indicated by asterisks and p-values. NS=not significant. (B-F) Neutralization and/or binding data for serum IgGs for recognition of (B) SARS-2 spike trimer, (C) SARS-2 RBD and SARS-2 pseudovirus, (D) SHC014 and WIV1 RBDs and corresponding pseudoviruses, (E) SARS RBD and SARS pseudovirus, (F) Yun 11, BM-4831, and BtKY72 RBDs.

Mice immunized with soluble SARS-2 S trimer (brown bars) showed no binding or neutralization except for autologous responses against SARS-2 after boosting (Fig. 3C-F). By contrast, sera from RBD-nanoparticle–immunized animals (red, green, yellow, and blue bars) exhibited binding to all RBDs (Fig. 3C-F; fig. S3A) and neutralization against all four strains after boosting (Fig. 3C-E), consistent with increased immunogenicities of multimerized antigen on nanoparticles versus soluble antigen (*27*, *28*). Homotypic SARS-2 nanoparticles, but not soluble SARS-2 trimer, induced heterologous responses to zoonotic RBDs and neutralization of heterologous coronaviruses (Fig. 3D-F). To address whether co-display of SARS-2 RBD along with other RBDs on mosaic-4a and mosaic-8 versus homotypic display of SARS-2 RBD (homotypic SARS-2) diminished anti-SARS-2 responses, we compared SARS-2–specific ELISA and neutralization titers for mosaic versus homotypic immunizations (Fig. 3C): there were no significant differences in IgG anti-SARS-2 titers for animals immunized with homotypic (red in Fig. 3C) versus mosaic nanoparticles (green and blue in Fig. 3C). Thus there was no advantage of immunization with a homotypic RBD-nanoparticle versus a mosaic-nanoparticle that included SARS-2 RBD in terms of the magnitude of immune responses against SARS-2.

We next compared serum responses against matched RBDs (RBDs present on an injected nanoparticle; gray horizontal shading) versus mismatched RBDs (RBDs not present on injected nanoparticle; red horizontal shading) (Fig. 3; fig. S3). Although SARS-2 RBD was not presented on mosaic-4b, antibody titers elicited by mosaic-4b immunization (yellow) were not significantly different than titers elicited by matched nanoparticle immunizations (homotypic SARS-2 (red), mosaic-4a (green), and mosaic-8 (blue)), and sera from boosted mosaic-4b–immunized mice neutralized SARS-2 pseudovirus (Fig. 3C). In other matched versus mismatched comparisons, sera showed binding and neutralization of SHC014 and WIV1 regardless of whether these RBDs were included on the injected nanoparticle (Fig. 3D), underscoring sharing of common epitopes among RBDs (Fig. 1A).

Demonstrating advantages of mosaic versus homotypic SARS-2 nanoparticles, sera from mosaic-8–immunized mice bound SHC014 and WIV1 RBDs significantly better after the prime than sera from homotypic SARS-2–immunized mice and retained better binding to SHC014 RBD after boosting (Fig. 3D). Thus the potential increased avidity of the homotypic SARS-2 nanoparticle displaying only one type of RBD over the mosaic-8 nanoparticles did not confer increased breadth. Moreover, mosaic-8–immunized and boosted sera were 7-44–fold more potent than sera from homotypic SARS-2–immunized animals in neutralizing SHC014 and WIV1 (Fig. 3D). Neutralization of the SHC014 and WIV1 pseudoviruses by mosaic-8 sera suggests that combining RBDs on a mosaic nanoparticle does not diminish the immune response against a particular RBD, also suggested by ELISA binding of sera to Rs4081 and RaTG13 (fig. S3A,B).

To further address whether RBD-nanoparticles elicited antibodies that recognized totally mismatched strains and SARS-CoV-2 RBD mutants, we evaluated sera for binding to SARS, Yun11, BM-4831, and BtKY72 RBDs (Fig. 3E,F), SARS-2 RBD mutants (fig. S3C), MERS-CoV RBD (fig. S3D), and for neutralization in SARS pseudovirus assays (Fig. 3E). We found no reductions in SARS-2 RBD binding as a result of mutations (Y453F, the “Danish mink variant” (*45*) or a Q493K/Q498Y/P499T triple mutant (*46*)) (fig. S3C), no binding of any elicited sera to MERS-CoV RBD (fig. S3D), and higher and more cross-reactive antibody responses for mosaic immunizations compared with homotypic SARS-2 immunizations: e.g., mosaic-8–primed and boosted animals showed significantly higher titers against SARS RBD than sera from homotypic SARS-2–immunized mice (Fig. 3E). After the prime, sera from the homotypic SARS-2–immunized animals did not neutralize SARS, whereas the mosaic-4b and mosaic-8 sera were neutralizing (Fig. 3E), perhaps facilitated by these nanoparticles including WIV1 RBD, which is related by 95% amino acid identity to SARS RBD (Fig. 1D). After boosting, SARS-2 and mosaic-4a sera were also neutralizing, although titers were ~4-fold lower than for mosaic-8–immunized animals (Fig. 3E). ELISA titers against other mismatched RBDs (Yun11, BM-4831, BtKY72) were significantly higher for sera collected after mosaic-8 priming compared to sera from homotypic SARS-2 priming, and heightened binding was retained after boosting (Fig. 3F). Thus mosaic nanoparticles, particularly mosaic-8, induce higher antibody titers against mismatched RBDs than homotypic SARS-2 nanoparticles, again favoring the co-display approach for inducing broader anti-coronavirus responses, especially after a single prime.

We investigated the potential for cross-reactive recognition using flow cytometry to ask whether B-cell receptors on IgG+ splenic B-cells from RBD-nanoparticle–boosted animals could simultaneously recognize RBDs from SARS-2 and Rs4081 (related by 70% sequence identity) (Fig. 1D; fig. S5). Whereas control animals were negative, all other groups showed B-cells that recognized SARS-2 and Rs4081 RBDs simultaneously, suggesting the existence of antibodies that cross-react with both RBDs (fig. S5E).

To compare antibodies elicited by RBD-nanoparticle immunization to antibodies elicited by SARS-CoV-2 infection, we repeated ELISAs against the RBD panel using IgGs from COVID-19 plasma donors (*47*) (Fig. 4). Most of the convalescent plasmas showed detectable binding to SARS-2 RBD (Fig. 4A). However, binding to other sarbecovirus RBDs (RaTG13, SHC014, WIV1, Rs4081 and BM-4831) was significantly weaker than binding to SARS 2 RBD, with many human plasma IgGs showing no binding above background (Fig. 4B-G). In addition, although convalescent plasma IgGs neutralized SARS-CoV-2 pseudoviruses, they showed weak or no neutralization of SARS, SHC014, or WIV1 pseudoviruses (Fig. 4H). These results are consistent with little to no cross-reactive recognition of RBDs from zoonotic coronavirus strains resulting from SARS-CoV-2 infection in humans.

**Figure 4.**
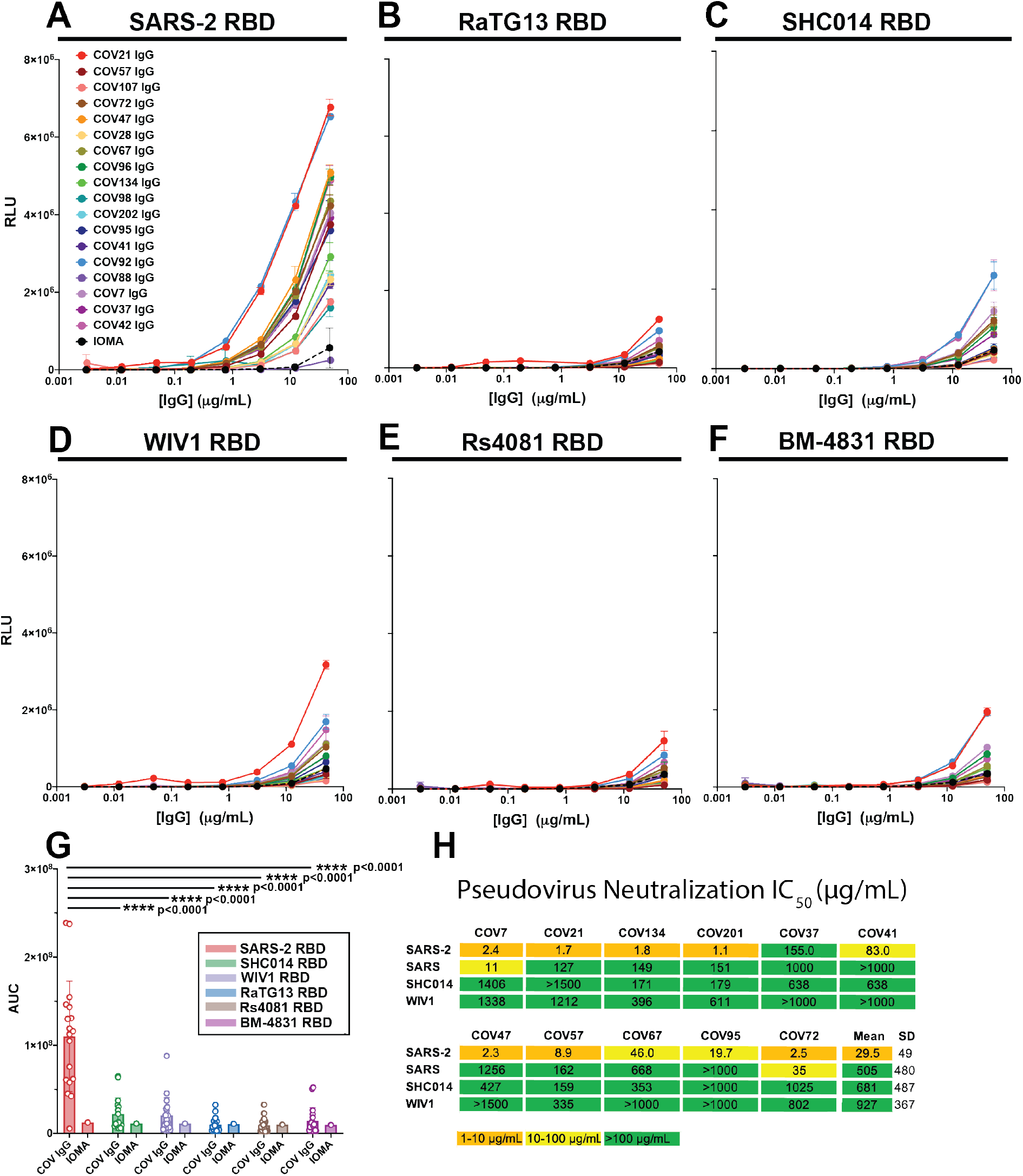
IgGs from convalescent COVID-19 plasma (*18*, *24*) show little to no cross-reactive responses. (A-F) Plasma IgG responses were evaluated by ELISA (data shown as binding curves with plasma names (*18*) listed) against RBDs from (A) SARS-2, (B) RaTG13, (C) SHC014, (D) WIV1, (E) Rs4081, and (F) BM-4831. Data points are plotted as the mean and standard deviation of duplicate measurements. IOMA, an anti-HIV-1 IgG (*50*), was used as a control. (G) ELISA results from panels A-F presented as area under the curve (AUC), where each dot represents one plasma sample, with means and standard deviations represented by rectangles (mean) and horizontal lines (SD). Significant differences between groups linked by horizontal lines are indicated by asterisks and p-values. (H) IC_50_ values for pseudotyped neutralization assays using IgGs from COV7, COV21, and COV72 plasmas (*18*) (evaluated at top concentrations of 1500 μg/mL) against the indicated strains. Mean=arithmetic mean IC_50_; SD=standard deviation.

In conclusion, we confirmed that multimerization of RBDs on nanoparticles enhances immunogenicity compared with soluble antigen (*33*, *48*) and further showed that homotypic SARS-2 nanoparticle immunization produced IgG responses that bound zoonotic RBDs and neutralized heterologous coronaviruses after boosting. By contrast, soluble SARS-2 S immunization and natural infection with SARS-CoV-2 resulted in weak or no heterologous responses in plasmas. Co-display of SARS-2 RBD along with diverse RBDs on mosaic nanoparticles showed no disadvantages for eliciting neutralizing antibodies against SARS-CoV-2 compared with homotypic SARS-2 nanoparticles, suggesting mosaic nanoparticles as a candidate vaccine to protect against COVID-19. Furthermore, compared with homotypic SARS-2 RBD particles, the mosaic co-display strategy demonstrated advantages for eliciting neutralizing antibodies against zoonotic sarbecoviruses, thus potentially also providing protection against emerging coronaviruses with human spillover potential., Neutralization of matched and mismatched strains was observed after mosaic priming, suggesting a single injection of a mosaic-RBD nanoparticle might be sufficient in a vaccine. Since COVID-19 convalescent plasmas showed little to no recognition of coronavirus RBDs other than SARS-CoV-2, COVD-19–induced immunity in humans may not protect against another emergent coronavirus. However, the mosaic nanoparticles described here could be used as described or easily adapted to present RBDs from newly-discovered zoonotic coronaviruses.

## Acknowledgements

We thank Karl Brune (Genie Biotech) for advice about mi3 production, Jesse Bloom (Fred Hutchinson) and Paul Bieniasz (Rockefeller University) for neutralization assay reagents, Jost Vielmetter and Caltech’s Beckman Institute Protein Expression Center for protein production, Andrew Flyak for help with flow cytometry, Marta Murphy for figures, COVID-19 plasma donors and Drs. Barry Coller and Sarah Schlesinger and the Rockefeller University Hospital Clinical Research Support Office and nursing staff, and Andrew Flyak and Andrew DeLaitsch for critical reading of the manuscript.

## Author contributions

A.A.C., C.O.B., and P.J.B. conceived and designed experiments. A.A.C., P.N.P.G., Y.E.L., P.R.H., S.O., and L.M.K. performed experiments, H-J.W. generated and validated SpyCatcher003-mi3, M.H. supervised the generation and validation of SpyCatcher003-mi3, A.A.C., J.R.K., A.P.W., C.O.B., M.C.N., and P.J.B. analyzed data and wrote the paper with contributions from other authors.

## Data and materials availability

All data are available in the main text of the supplementary materials. Materials are available upon request to with a signed Material Transfer Agreement.

## Supplementary content

Materials and Methods, Figs. S1 to S5, References (*51*–*58*).

## Materials and Methods

### Phylogenetic tree

A sequence alignment of coronavirus RBD domains was made using Clustal Omega (*51*). A phylogenetic tree was calculated from this amino acid alignment using PhyML 3.0 (*52*), and a figure of this tree was made using PRESTO (http://www.atgc-montpellier.fr/presto).

### Expression of RBD and S proteins

Mammalian expression vectors encoding the RBDs of SARS-CoV-2 (GenBank MN985325.1; S protein residues 319-539) and SARS-CoV S (GenBank AAP13441.1; residues 318-510) with an N-terminal human IL-2 or Mu phosphatase signal peptide were previously described (*47*). Expression vectors were constructed similarly for RBDs from the following other sarbecovirus strains: RaTG13-CoV (GenBank QHR63300; S protein residues 319-541), SHC014-CoV (GenBank KC881005; residues 307-524), Rs4081-CoV (GenBank KY417143; S protein residues 310-515), pangolin17-CoV (GenBank QIA48632; residues 317-539), RmYN02-CoV (GSAID EPI_ISL_412977; residues 298-503), Rf1-CoV (GenBank DQ412042; residues 310-515), W1V1-CoV (GenBank KF367457; residues 307-528), Yun11-CoV (GenBank JX993988; residues 310-515), BM-4831-CoV (GenBank NC014470; residues 310-530), BtkY72-CoV (GenBank KY352407; residues 309-530). Two versions of each RBD expression vector were made: one including a C-terminal hexahistidine tag (G-HHHHHH) and SpyTag003 (RGVPHIVMVDAYKRYK) (*43*) (for coupling to SpyCatcher003-mi3) and one with only a hexahistidine tag (for ELISAs). Biotinylated SARS-CoV-2 and Rs4081 RBDs were produced by cotransfection of Avi/His-tagged RBD expression plasmids with an expression plasmid encoding an ER-directed BirA enzyme (kind gift of Michael Anaya, Caltech). RBD proteins were purified from transiently-transfected Expi293F cell (Gibco) supernatants by nickel affinity and size-exclusion chromatography (*47*). Peak fractions corresponding to RBDs were identified by SDS-PAGE and then pooled and stored at 4°C. A trimeric SARS-CoV-2 ectodomain with 6P stabilizing mutations (*53*) was expressed and purified as described (*24*). Correct folding of the soluble SARS-CoV-2 S trimer was verified by a 3.3 Å cryo-EM structure of a neutralizing antibody complexed with the trimer preparation used for immunizations (*24*). To prepare fluorochrome-conjugated streptavidin-tetramerized RBDs, biotinylated SARS-2 and Rs4081 RBDs were incubated with streptavidin-APC (eBioscience™) and streptavidin-PE (ThermoFisher), respectively, overnight at 4°C at a 1:1 molar ratio of RBD to streptavidin subunit.

### Preparation of human plasma IgGs

Plasma samples collected from COVID-19 convalescent and healthy donors are described in (*18*). Human IgGs were isolated from heat-inactivated plasma samples using 5-mL HiTrap MabSelect SuRe columns (GE Healthcare Life Sciences) as described (*24*).

### Preparation of RBD-mi3 nanoparticles

SpyCatcher003-mi3 particles were prepared by purification from BL21 (DE3)-RIPL *E coli* (Agilent) transformed with a pET28a SpyCatcher003-mi3 gene (including an N-terminal 6x-His tag) as described (*54*). Briefly, cell pellets from transformed bacterial were lysed with a cell disruptor in the presence of 2.0 mM PMSF (Sigma). Lysates were spun at 21,000xg for 30 min, filtered with a 0.2 μm filter, and mi3 particles were isolated by Ni-NTA chromatography using a pre-packed HisTrap™ HP column (GE Healthcare). Eluted particles were concentrated using an Amicon Ultra 15 mL 30K concentrator (MilliporeSigma) and purified by SEC using a HiLoad^®^ 16/600 Superdex^®^ 200 (GE Healthcare) column equilibrated with 25 mM Tris-HCl pH 8.0, 150 mM NaCl, 0.02% NaN3 (TBS). SpyCatcher-mi3 particles were stored at 4°C and used for conjugations for up to 1 month after filtering with a 0.2 μm filter or spinning at 21,000xg for 10 min.

Purified SpyCatcher003-mi3 was incubated with a 3-fold molar excess (RBD to mi3 subunit) of purified SpyTagged RBD (either a single RBD for making homotypic SARS-CoV-2 RBD particles or an equimolar mixture of four or eight RBDs for making mosaic particles) overnight at room temperature in TBS. Conjugated mi3 particle were separated from free RBDs by SEC on a Superose 6 10/300 column (GE Healthcare) equilibrated with PBS (20 mM sodium phosphate pH 7.5, 150 mM NaCl). Fractions corresponding to conjugated mi3 particles were collected and analyzed by SDS-PAGE. Concentrations of conjugated mi3 particles were determined using a Bio-Rad Protein Assay.

### Immunizations

Animal procedures and experiments were performed according to protocols approved by the IACUC. Experiments were done using 4-6 week old female Balb/c mice (Charles River Laboratories), with 5 animals each for cohorts immunized with soluble SARS-CoV-2 S or SpyCatcher003-mi3, and 10 animals each for remaining cohorts (Fig 3A). Immunizations were carried out with intraperitoneal (ip) injections of either 5 μg of conjugated RBD (calculated as the mass of the RBD, assuming 100% efficiency of conjugation to SpyCatcher003-mi3), 5 μg of soluble SARS-CoV-2 S, or 6 μg of unconjugated SpyCatcher003-mi3, in 100 μL of 50% v/v AddaVax™ adjuvant (Invivogen). Animals were boosted 4 weeks after the prime with the same quantity of antigen in adjuvant. Animals were bled every 2 weeks via tail veins, and then euthanized 8 weeks after the prime (Day 56, 57) and bled through cardiac puncture. Blood samples were allowed to clot at room temperature in MiniCollect^®^ Serum and Plasma Tubes (Greiner), and serum was harvested, preserved in liquid nitrogen, and stored at −80°C until use.

Sera for ELISAs were collected at Day 14 (Prime) and Day 42 (Boost). Sera for neutralization assays were collected at Day 28 (Prime) and Day 56 (Boost) (Fig. 3, fig. S3).

### ELISAs

10 μg/ml of a purified RBD (not SpyTagged) in 0.1 M NaHCO_3_ pH 9.8 was coated onto Nunc^®^ MaxiSorp™ 384-well plates (Sigma) and stored overnight at 4°C. Plates were washed with Tris-buffered saline with 0.1% Tween 20 (TBS-T) after blocking with 3% bovine serum albumin (BSA) in TBS-T for 1 hr at room temperature. Mouse serum was diluted 1:100 and then serially diluted by 4-fold with TBS-T/3% BSA and added to plates for 3 hr at room temperature. A 1:50,000 dilution of secondary HRP-conjugated goat anti-mouse IgG (Abcam) was added after washing for 1 hr at room temperature. Plates were developed using SuperSignal™ ELISA Femto Maximum Sensitivity Substrate (ThermoFisher) and read at 425 nm. Curves were plotted and integrated to obtain the area under the curve (AUC) using Graphpad Prism 8.3 assuming a one-site binding model with a Hill coefficient (Fig. 3; fig. S3). We also calculated EC_50_s and endpoint titers, which were determined using the dilution that was at or below the mean + 2 x the standard deviation of the plate control (no primary serum added) for ELISA binding data (fig. S3E,F). AUC calculations were used as they better capture changes in maximum binding (*55*). Statistical significance of titer differences between groups were calculated using Tukey’s multiple comparison test using Graphpad Prism 8.3.

### Neutralization assays

SARS-CoV-2, SARS, WIV1, and SHC014 pseudoviruses based on HIV lentiviral particles were prepared as described (*18*, *56*) using genes encoding S protein sequences lacking C-terminal residues in the cytoplasmic tail: 21 amino acid deletions for SARS-CoV-2, WIV1, and SHC014 and a 19 amino acid deletion for SARS-CoV. IC_50_ values derived from this pseudotyped neutralization assay method were shown to quantitatively correlate with results from neutralization assays using authentic SARS-CoV-2 virus (*44*). For pseudovirus neutralization assays, four-fold serially diluted sera from immunized mice were incubated with a pseudotyped virus for 1 hour at 37°C. After incubation with 293T_ACE2_ target cells for 48 hours at 37°C, cells were washed twice with phosphate-buffered saline (PBS) and lysed with Luciferase Cell Culture Lysis 5x reagent (Promega). NanoLuc Luciferase activity in lysates was measured using the Nano-Glo Luciferase Assay System (Promega). Relative luminescence units (RLUs) were normalized to values derived from cells infected with pseudotyped virus in the absence of serum. Half-maximal inhibitory dilutions (ID_50_ values) were determined using 4-parameter nonlinear regression in AntibodyDatabase (*57*). Statistical significance of titer differences between groups were calculated using Tukey’s multiple comparison test of ID_50_s converted to log^10^ scale using Graphpad Prism 8.3.

### Statistical Analysis

Comparisons between groups for ELISAs and neutralization assays were calculated with one-way analysis of variance (ANOVA) using Tukey’s post hoc test in Prism 9.0 (Graphpad). For correlation analysis between ELISA and neutralization titers, significance (p), Spearman coefficients (r_s_), and linear plots were calculated using Prism 9.0 (Graphpad). Differences were considered significant when p values were less than 0.05. Exact p values are in relevant figure near each corresponding line, with asterisks denoting level of significance (* denotes 0.01<p<0.05, ** denotes 0.001<p<0.01, *** denotes 0.0001<p<0.001, and **** denotes p<0.0001).

### Flow cytometry

B-cell analysis using flow cytometry was carried out as described (*54*). Briefly, single-cell suspensions were prepared from mouse spleens using mechanical dissociation, and red blood cells were removed using ACK lysing buffer (Gibco). The white blood cell preparation was enriched for IgG+ B-cells using the negative selection protocol in a mouse memory B-cell isolation kit (Miltenyi). The following commercial reagents were used to stain enriched splenocytes: CD4-APC-eFluor 780 (clone: RM4-5), F4/80-APC-eFluor 780 (clone: BM8), CD8a-APC-eFluor 780 (clone: 53-6.7), Ly-6G-APC-eFluor 780 (clone: RB6-8C5), IgM-APC-eFluor 780 (clone: II/41) (Thermo Fisher Scientific), CD19-FITC (clone: 6D5) (Biolegend), IgG1 BV421 (clone: X40) and IgG2 BV421 (clone: R19-15) (BD Bioscience). SARS-2 RBD-APC and Rs4081 RBD-PE for used to identify antigen-specific B-cells. Cell viability was analyzed with Fixable Viability Stain 700 (BD Bioscience). Stained cells were analyzed with a SY3200 Cell Sorter (Sony) configured to detect 6 fluorochromes. 2,000,000 events were collected per sample and analyzed via FlowJo software (TreeStar).

---

**Fig. S1.**
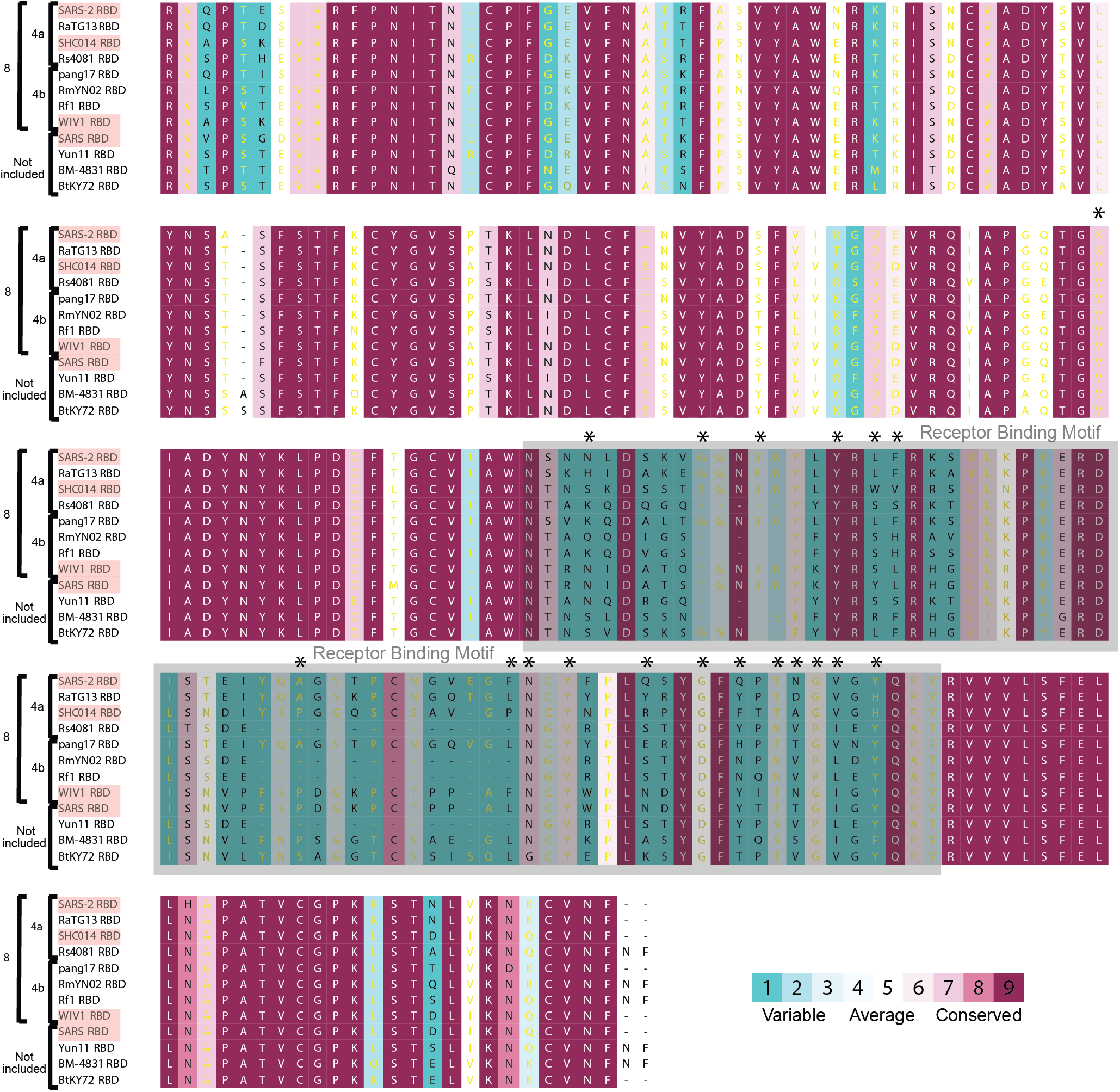
Alignment of RBD sequences used for making mosaic particles. Sequences shown are for the RBDs of SARS-CoV-2 (SARS-2, GenBank: MN985325.1), RaTG13 (QHR63300), SHC014 (RsSHC014, KC881005), Rs4081 (KY417143), PCoV_GX-P5L (pang17) (QIA48632), RmYN02 (GSAID EPI_ISL_412977), Rf1 (DQ412042), WIV1 (KF367457), SARS-CoV (AAP13441.1), Yun11 (Cp/Yunnan2011, JX993988), BM-4831 (BM48-31/BGR/2008, NC014470), and BtKY72 (KY352407). SARS-2 RBD residues that interact directly with ACE2 (*58*) are indicated by an asterisk. We note that antibody neutralization by direct binding of ACE2-binding residues does not represent the only mechanism of neutralization for ACE2-tropic viruses. This has been shown for monoclonal human antibodies derived from COVID-19 patients: some neutralizing antibodies do not directly interact with the ACE2-binding site on RBD (for example, class 3 anti-SARS-CoV-2 neutralizing antibodies (*24*)). Red shading indicates strains known to use ACE2 as a receptor.

**Fig. S2.**
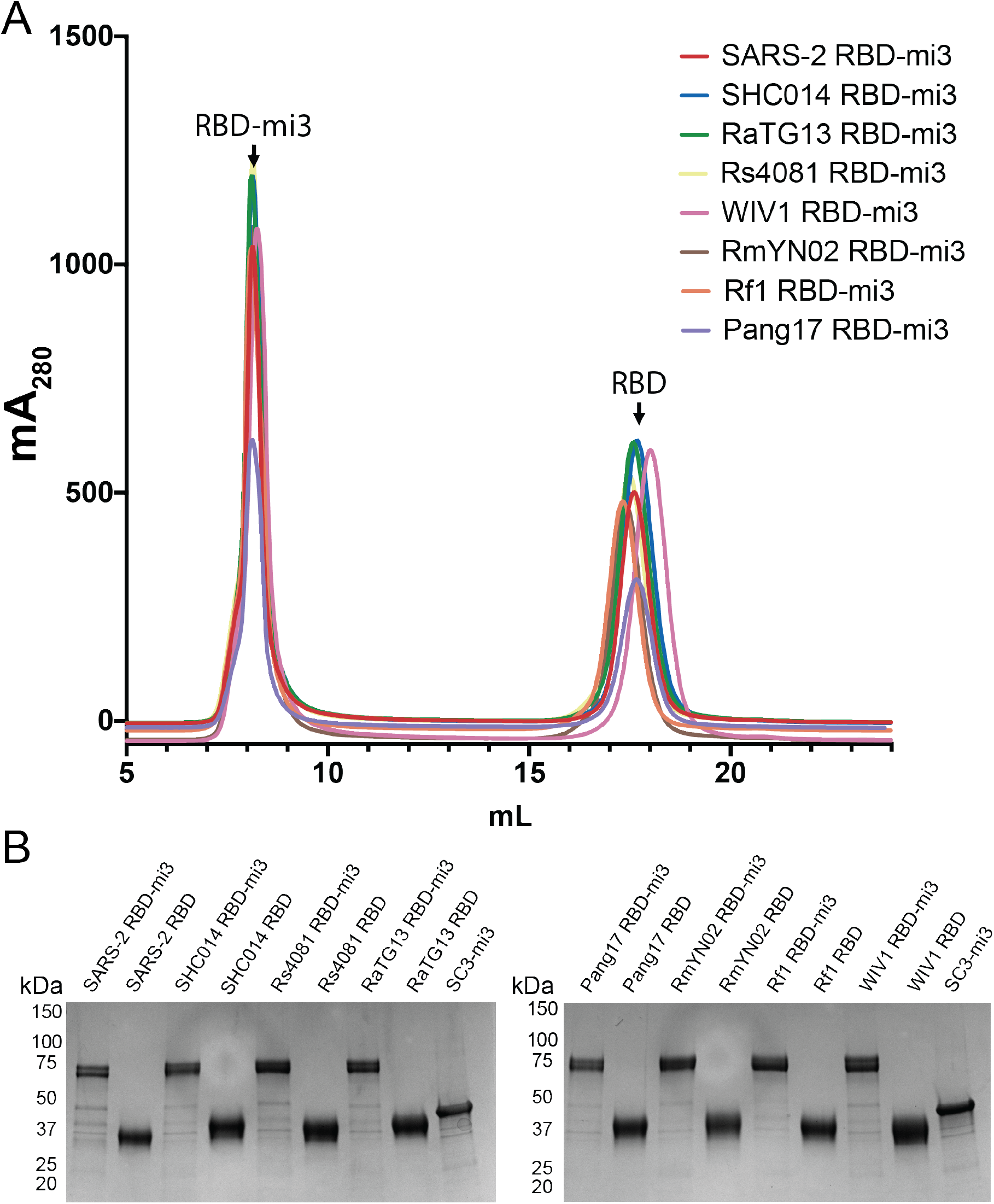
RBDs from the eight sarbecovirus S proteins conjugate equivalently to SpyCatcher003-mi3, suggesting a statistical mixture of RBDs on mosaic particles. (A) SEC profiles showing separation of RBD nanoparticles and free RBD proteins. (B) Coomassie-stained SDS-PAGE of RBD-coupled nanoparticles, free RBD proteins, and uncoupled SpyCatcher003-mi3 particles (SC3-mi3).

**Fig. S3.**
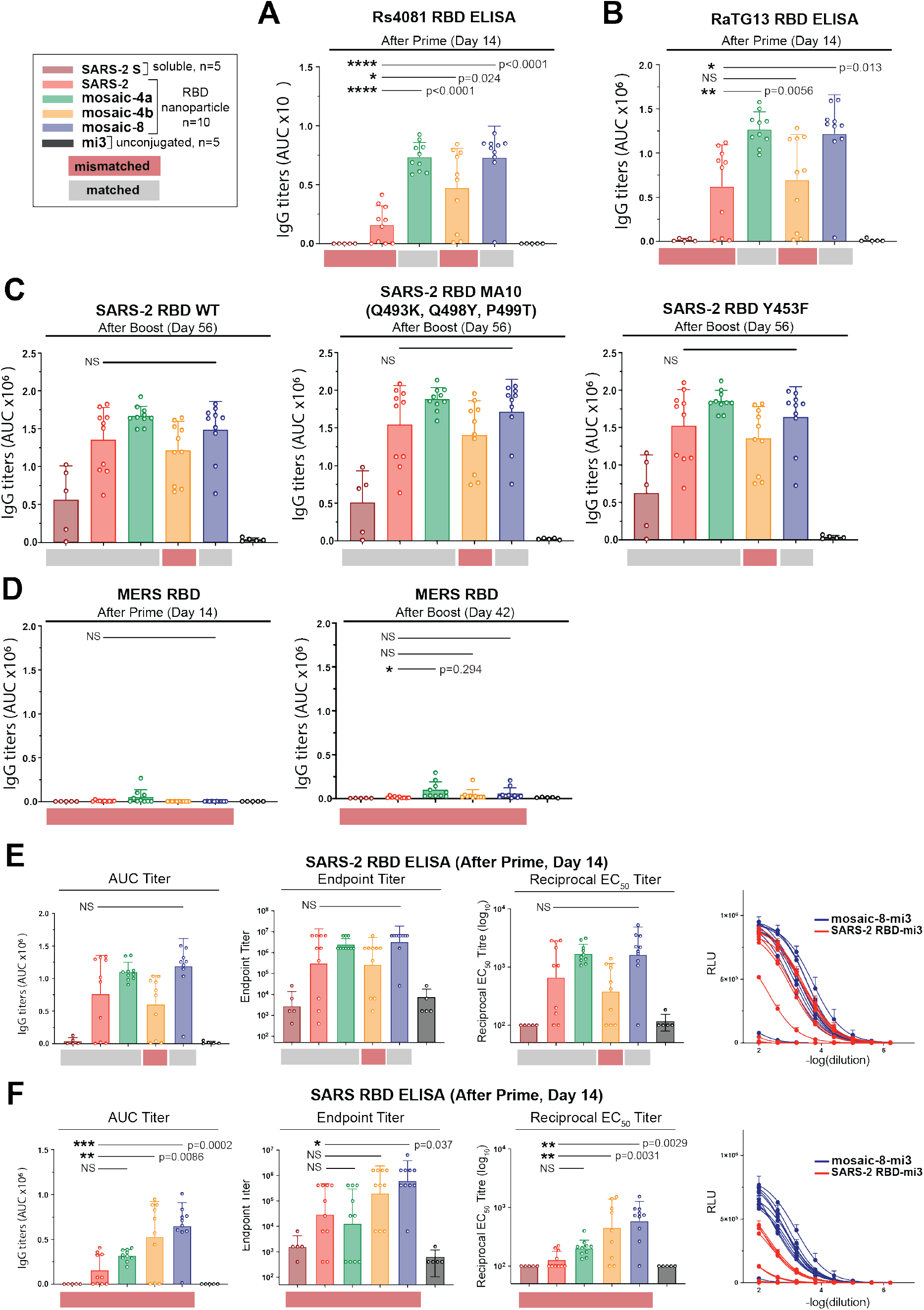
Day 14 serum IgG responses to RBDs evaluated by ELISA shown as area under the curve (AUC) from mice immunized with soluble SARS-CoV-2 S trimers (SARS-2 S) or RBDs on nanoparticles (homotypic SARS-2, mosaic-4a, mosaic-4b, mosaic-8, or unconjugated SpyCatcher003-mi3 (mi3)). Each dot represents serum from one animal, with means and standard deviations represented by rectangles (mean) and horizontal lines (SD). RBDs from strains that were not present on an immunized particle or were present on an immunized particle are indicated by red and gray rectangles, respectively, below the ELISA data. Significant differences between groups linked by horizontal lines are indicated by asterisks and p-values. NS=not significant. (A,B) Binding of serum IgGs to (A) Rs4081 and (B) RaTG13 RBDs. (C) Binding of serum IgGs to SARS-2 RBD (left), a triple RBD mutant in a mouse-adapted SARS-CoV-2 (*46*) that includes substitutions adjacent to the N501Y RBD mutation in an emergent UK SARS-CoV-2 lineage (https://virological.org/t/preliminary-genomic-characterisation-of-an-emergent-sars-cov-2-lineage-in-the-uk-defined-by-a-novel-set-of-spike-mutations/563) (middle), and Y453F, the “Danish mink variant” (*45*) (right). (D) Binding of serum IgGs to RBD from MERS-CoV (a non-ACE2-binding merbecovirus, representing a different subgenus from sarbecoviruses). (E,F) Comparison of ELISA data for serum binding to selected RBDs presented as AUC, endpoint titers, midpoint titers, or binding curves. Day 14 serum IgG responses to (E) SARS-2 or (F) SARS RBDs evaluated by ELISA shown as AUC (left), endpoint titers (middle left), midpoint (EC_50_) titers (middle right), or binding curves (right). For AUC, each dot represents serum from one animal, with means and standard deviations represented by rectangles (mean) and horizontal lines (SD). For endpoint and midpoint titers, each dot represents serum from one animal, with geometric means and geometric standard deviations represented by rectangles (mean) and horizontal lines (SD). Binding curves are shown with data points representing the mean and SD of duplicate measurements fit to a binding model (see Methods) for animals immunized with mosaic-8 and homotypic SARS-2.

**Fig. S4.**
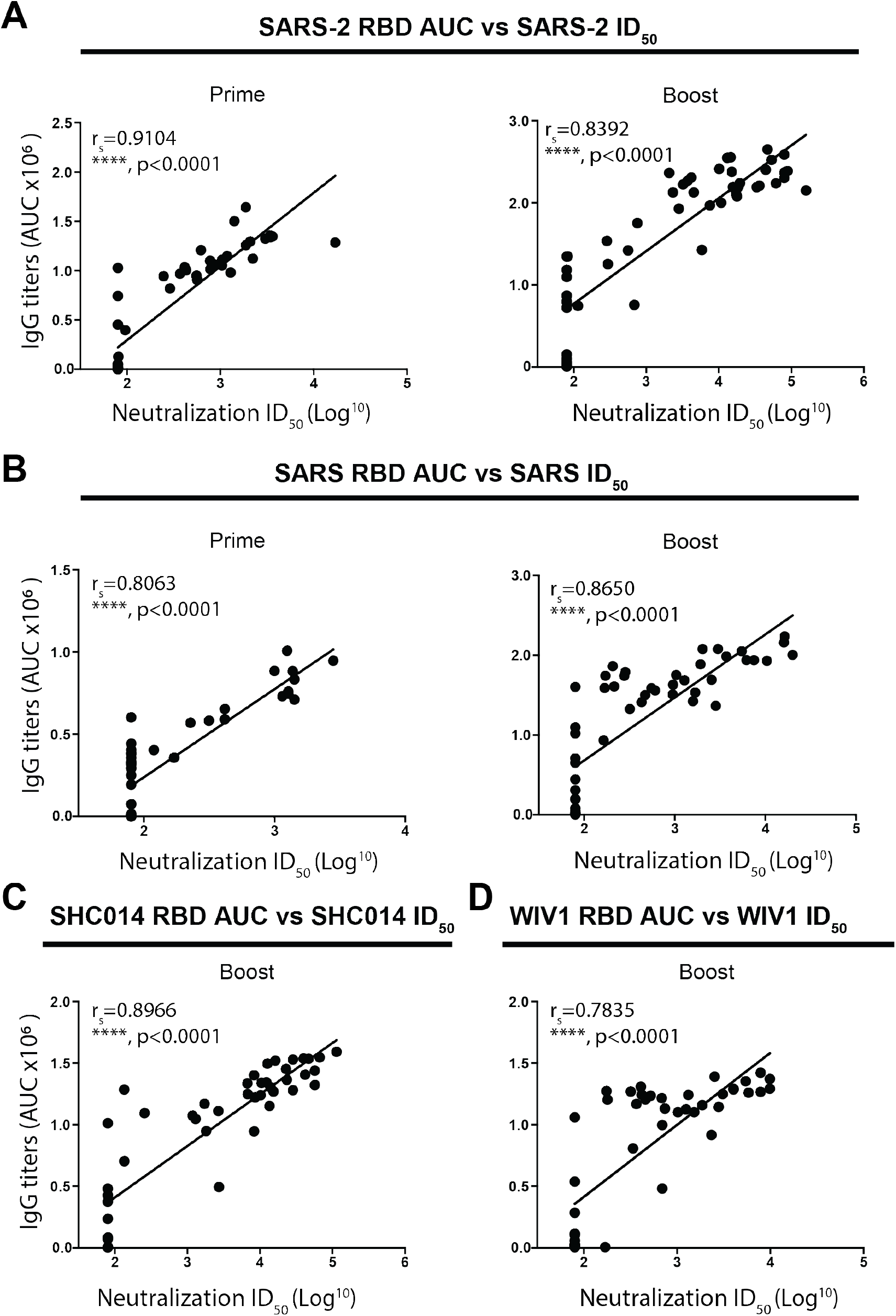
Correlation of ELISA and neutralization titers. Spearman correlation coefficients (r_S_) and p-values shown for graphs of anti-RBD ELISA titers (AUC) versus pseudovirus neutralization ID_50_ values; significance indicated as asterisks. (A) SARS-2. (B) SARS. (C) SHC014. (D) WIV1.

**Fig S5.**
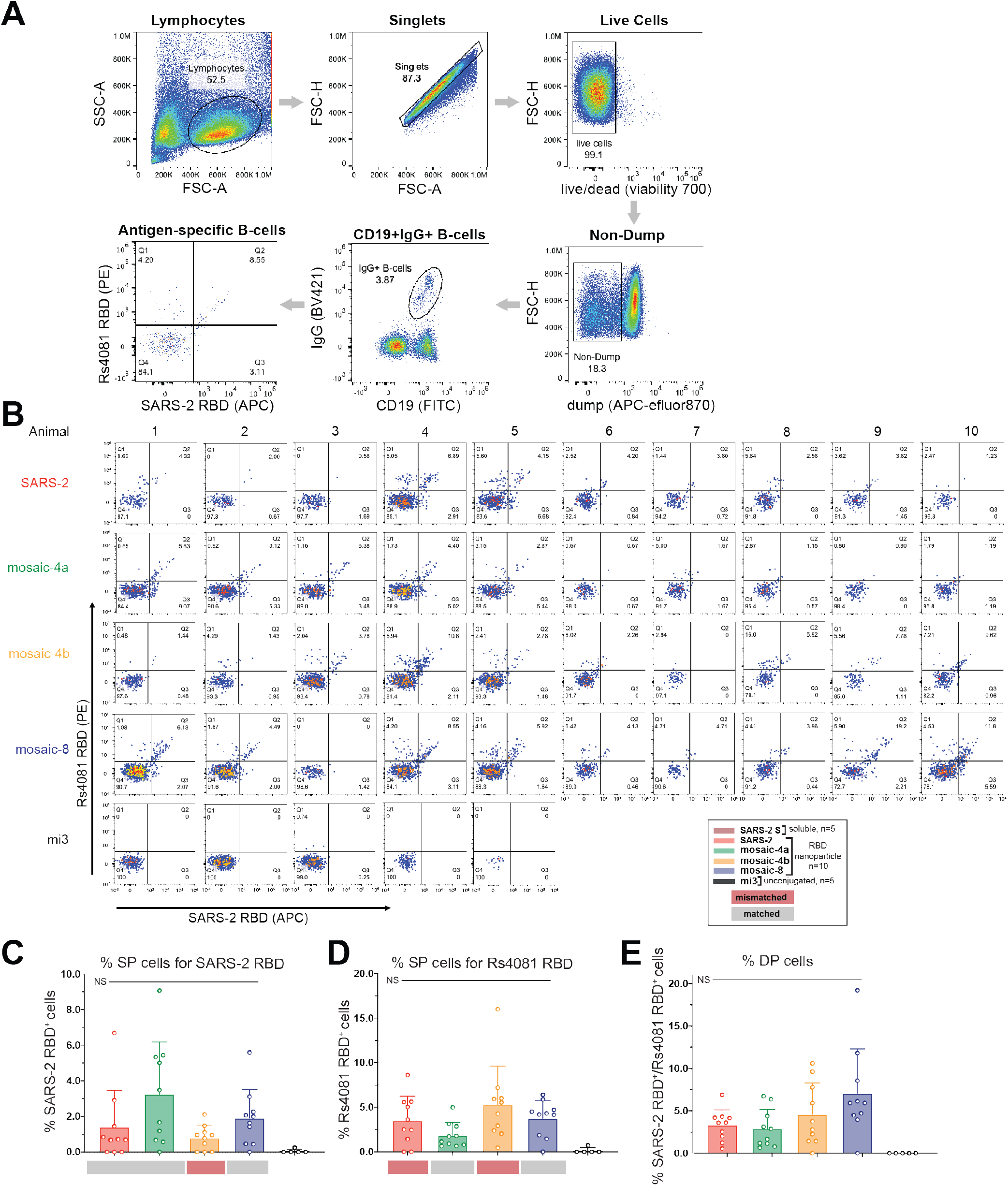
Antigen-specific IgG^+^ B-cell analysis of splenocytes isolated from animals immunized with mosaic-RBD nanoparticles. (A) Flow cytometry gating strategy for characterizing RBD-specific IgG^+^ B-cells isolated from splenocytes. Anti-CD4, anti-CD8, anti-F4/80, anti-Ly6G, and anti-IgM were used in the dump to remove T-cells, macrophages, and IgM^+^ B-cells. Antigenspecific IgG^+^ B-cells were isolated using labeled anti-CD19 and anti-IgG antibodies, and probed for binding RBD with a pair of fluorophore-conjugated RBD tetramers (SARS-2 RBD and Rs4081 RBD). (B) Complete flow cytometry analysis for antigen-specific IgG^+^ splenocytes isolated from animals immunized with mosaic-RBD particles. The 4-way gate shown for each animal separates each population of RBD single-positive and double-positive cells and was used for the % antigen-specific populations shown in panels C-E. Q1 represents the Rs4081 RBD^+^ population, Q2 represents the Rs4081 RBD^+^ / SARS-2 RBD^+^ population, Q3 represents the SARS-2 RBD^+^ population, and Q4 represents the RBD^-^ population. (C-E) Percent single-positive (SP) and double-positive (DP) cells for the indicated groups. Significant differences between groups linked by horizontal lines are indicated by asterisks and p-values. NS = not significant. (C) Percent SARS-2 RBD^+^ B-cells within the IgG^+^ B-cell population. (D) Percent Rs4081 RBD^+^ B-cells within the IgG^+^ B-cell population. (E) Percent SARS-2 RBD^+^ / Rs4081 RBD^+^ B-cells within the IgG^+^ B-cell population.

